# petiteFinder: An automated computer vision tool to compute Petite colony frequencies in baker’s yeast

**DOI:** 10.1101/2022.05.12.491699

**Authors:** Christopher J. Nunn, Eugene Klyshko

## Abstract

Mitochondrial respiration is central to cellular and organismal health in eukaryotes. In baker’s yeast, however, respiration is dispensable under fermentation conditions. Because yeast are tolerant of this mitochondrial dysfunction, yeast are widely used by biologists as a model organism to ask a variety of questions about the integrity of mitochondrial respiration. Fortunately, baker’s yeast also display a visually identifiable Petite colony phenotype that indicates when cells are incapable of respiration. Petite colonies are smaller than their Grande (wild-type) counterparts, and their frequency can be used to infer the integrity of mitochondrial respiration in populations of cells. In this study, we introduce a deep learning enabled tool, *petiteFinde*r, to leverage the Petite colony phenotype and increase the throughput of the Petite frequency assay. This automated computer vision tool detects Grande and Petite colonies and computes Petite colony frequencies from scanned images of Petri dishes. It addresses issues in scalability and reproducibility of the Petite colony assay which currently relies on laborious manual colony counting methods. Combined with the detailed experimental protocols we provide, we believe this study can serve as a foundation to standardize this assay. Finally, we comment on how Petite colony detection as a computer vision problem highlights ongoing difficulties with small object detection in existing object detection architectures.

## Introduction

Eukaryotic cells have numerous mitochondria containing multiple copies of their genome, mitochondrial DNA (mtDNA). In most eukaryotes, mtDNA encodes a subset of proteins involved in oxidative phosphorylation, while the remainder is encoded in nuclear DNA (nDNA). Saccharomyces cerevisiae, or baker’s yeast, is a widely used model organism to study the interdependence of mtDNA and nDNA. This is largely due to the dispensability of respiration in yeast under fermentable conditions, which allows cells to propagate without mitochondrial/nuclear-encoded genes involved in respiration. In particular, tolerance to mutations in mtDNA, which would otherwise be fatal to cells in other model systems, enables the study of mtDNA dynamics in yeast, including interesting questions regarding mtDNA maintenance, selection, and mutation.

Since the discovery of the Petite colony phenotype in yeast, this phenotype has played a central role in studying the interplay of nDNA and mtDNA (Ephrussi et al., 1949; 1953). As the name suggests, the Petite phenotype is characterized by small, more translucent colonies compared to Grande (wild-type) colonies under fermentable conditions. These colonies are smaller and have lower cell density due to the inability to switch to respiration once fermentable carbon sources have been consumed as the colony spreads on an agar surface. The ease of distinguishing Petite from Grande colonies visually, with nothing but a Petri dish and its agar surface, is part of what has made the Petite phenotype popular as an indicator of mitochondrial function.

Measurements of Petite colony frequencies have found various uses in the literature. By transplanting mtDNA into yeast strains with various genetic backgrounds, changes in Petite frequencies resulting from this transplantation have been used to understand what effect variants in nuclear genes and mtDNA structures have on mtDNA stability (Contamine & Picard, 2000, Dimitrov et al., 2009). Petite frequencies have also been shown to be useful as a screen to determine what nuclear genes are responsible for mtDNA recombination that affect mtDNA integrity (Ling & Shibata, 2002, 2004; Ling et al., 2007; Ling et al., 2019), and chemicals that induce mitochondrial maintenance, or mitophagy (Karavaeva et al., 2017). Petite frequency measurements have been integrated into a powerful approach that discovered numerous genes involved in mitochondrial biogenesis (Hess et al., 2009). Furthermore, mating cells with wild-type and mutated mtDNAs and measuring the fraction of progeny that themselves are Petite provides access to the selection and dynamics of competing mtDNAs (de Zamaroczy et al., 1981; Nunn & Goyal, 2022).

While the Petite frequency assay is both accessible and versatile, current methods of Petite colony identification are laborious. These often involve manual annotation of Petri dishes under growth conditions where Petite/Grande colony differences are perceptible or replica plating onto non-fermentable media (see for example the methods in Dimitrov et al., 2009; Hess et al., 2009; Karavaeva et al., 2017). Annotation under different growth conditions across laboratories, which can influence Petite vs. Grande colony characteristics, also hinders reproducibility. To address these issues in scalability and reproducibility, which limits the statistical power of this assay, in this study we develop an automated computer vision tool to detect Petite/Grande colonies and determine Petite frequencies from scanned images of Petri dishes. Our tool, *petiteFinder*, is built on the open-source object detection toolbox MMDetection (Wang et al., 2019) and a computer vision library SAHI (Akyon et al., 2022), which improves small object detection via image slicing. It uses a Faster R-CNN (Ren et al., 2016) object detection architecture built on top of a ResNet-50 backbone (He et al., 2015) coupled with a feature pyramid network (FPN) (Lin et al., 2017). Beyond colony detection, *petiteFinder* also enables users to amend colony classification predictions with a user-friendly interface.

## Methods

### Acquisition of the Grande/Petite colony dataset

To generate a dataset for this tool, we performed plating experiments where yeast colonies were grown on an agar surface in Petri dishes. We conducted these experiments with fermentable media (both synthetic and complete) that can be constructed to support baker’s yeast strains with any auxotrophies (see details in *Media and growth conditions*). The only constraint on the media was a carbon composition of 3% glycerol and 0.1% glucose that allows for robust visual identification of Grande/Petite colonies after 3-5 days of growth on agar at 30°C (Dimitrov et al., 2009). With this carbon composition, Petite colonies appear smaller and more translucent than their Grande counterparts on an agar surface. On synthetic media (SC-ura-trp), we mated a Grande strain and a variety of Petite strains. We plated this mixture to identify whether or not the progeny of mated parents were Petite or Grande. On complete media (YPADG), we plated cells from a single haploid strain to identify spontaneous transitions from Grande to Petite cells. These experiments yielded Petri dishes with Petite colony frequencies ranging from ∼0 to 0.9.

Following these experiments, Petri dishes were scanned with an image scanner, six at a time (see *Petri dish imaging*, Figure 1a). Images were then cropped to contain an individual petri dish with the aid of a 3D printed scanner insert that fixed their positions and reduced refraction due to adjacent plates. These images were manually annotated using the LabelMe package (Russell et al., 2008), where bounding boxes were drawn around Grande and Petite colonies and assigned to their corresponding labels by an expert in Petite/Grande colony identification. The resulting labeled dataset is a collection of 83 petri dish images, 59 in synthetic media and 24 in complete media. There are 5799 bounding box annotations encompassing colonies, with 2716 Petite and 3083 Grande labels in total. Variation in agar concentrations within synthetic media poured into plates in this dataset produced 20 images we consider “non-ideal” due to the diffuseness of colonies, and 39 images we consider “ideal” (Figure 1b). Images of complete media were largely consistent and did not exhibit this variability. All types of images were used as training data. This experimental variation, both at the level of media and the types of experiments that generated these plates, is further augmented during training to produce a robust computer vision model. Images were split into 60% training, 20% validation, and 20% test sets.

**Figure 1:**
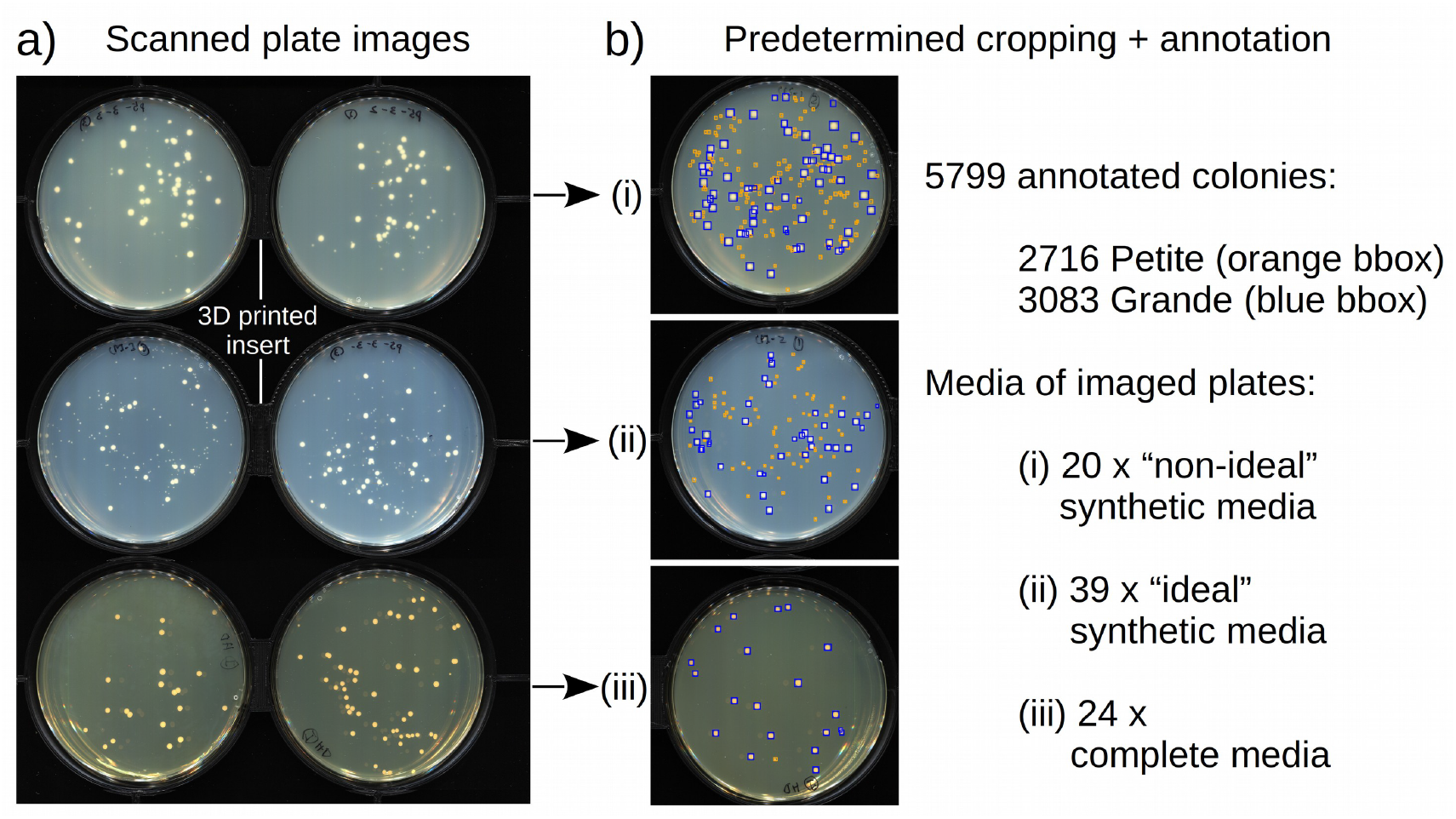
An overview of the labeled dataset. a) A composite image showing Petri dishes containing synthetic (top 4 Petri dishes) and complete media (bottom 2 Petri dishes) with yeast colonies on their surface. Six petri dishes at a time were placed in a 3D printed insert (black structure bordering petri dish images) and scanned on a computer scanner bottom-up. The synthetic media was SC-ura-trp and the complete media was YPADG (both 0.1% glucose and 3% glycerol carbon source). b) Individual plate images were cropped out of the large image, and 83 of these images were annotated using the LabelMe annotation tool (Russell et al., 2008). Grande and Petite colonies are indicated by blue and orange bounding boxes, respectively. Experimental variation in media preparation/pouring of plates produces diffuse low-quality scans (non-ideal) and sharper plate images (ideal) as shown by example plate crops (i) and (ii) for synthetic media. Plate crop (iii) is complete media that has uniform agar opacity across all images.

### Model architecture

The detection and classification of colonies is an object detection problem that is commonly solved in computer vision through convolutional neural networks (CNN). Object detection frameworks built on region-based convolutional neural networks (RCNNs) remain among top performers in object detection competitions (Xiao et al., 2020). However, small object detection is challenging for most RCNN approaches, where lower-level features (in early layers of the network) of small objects are lost through coarse-graining via convolutions and pooling. The Faster-RCNN architecture (Ren et al., 2016) coupled with a feature pyramid network (FPN) has been shown to improve performance in small object detection (Lin et al., 2017). Specifically, the FPN achieves high accuracy detection of small objects by propagating high-level features down to lower layers in the network and sharing feature maps of multiple scales with the region proposal component of the detector.

Despite adopting the architectural advantages of FPN for small object detection, a Faster-RCNN ResNet-50 FPN pipeline performed poorly in Petite colony detection on intact images of Petri dishes. We attributed this to the initial downsampling of images (1024×1024) we performed to fit models on consumer GPUs. This downsampling can eliminate Petite colonies in images due to their small size. To address this poor performance, we employed SAHI (Akyon et al., 2022), a computer vision package that performs model inference through image slicing. Instead of intact images, small slices of images are passed through the Faster-RCNN ResNet-50 FPN architecture, which increases the relative size of colonies to the background in the network. Following object detection on all image slices from sliding windows over the entire image, slice predictions are merged in a greedy non-maximal merging algorithm (NMM). This algorithm operates like non-maximal suppression (NMS), but instead of eliminating bounding boxes, it simply merges them above a particular intersection over smaller area (IOS) threshold. The final output of the model operating on an image is a collection of predicted colony bounding boxes with Grande/Petite classification probability (Figure 2).

**Figure 2:**
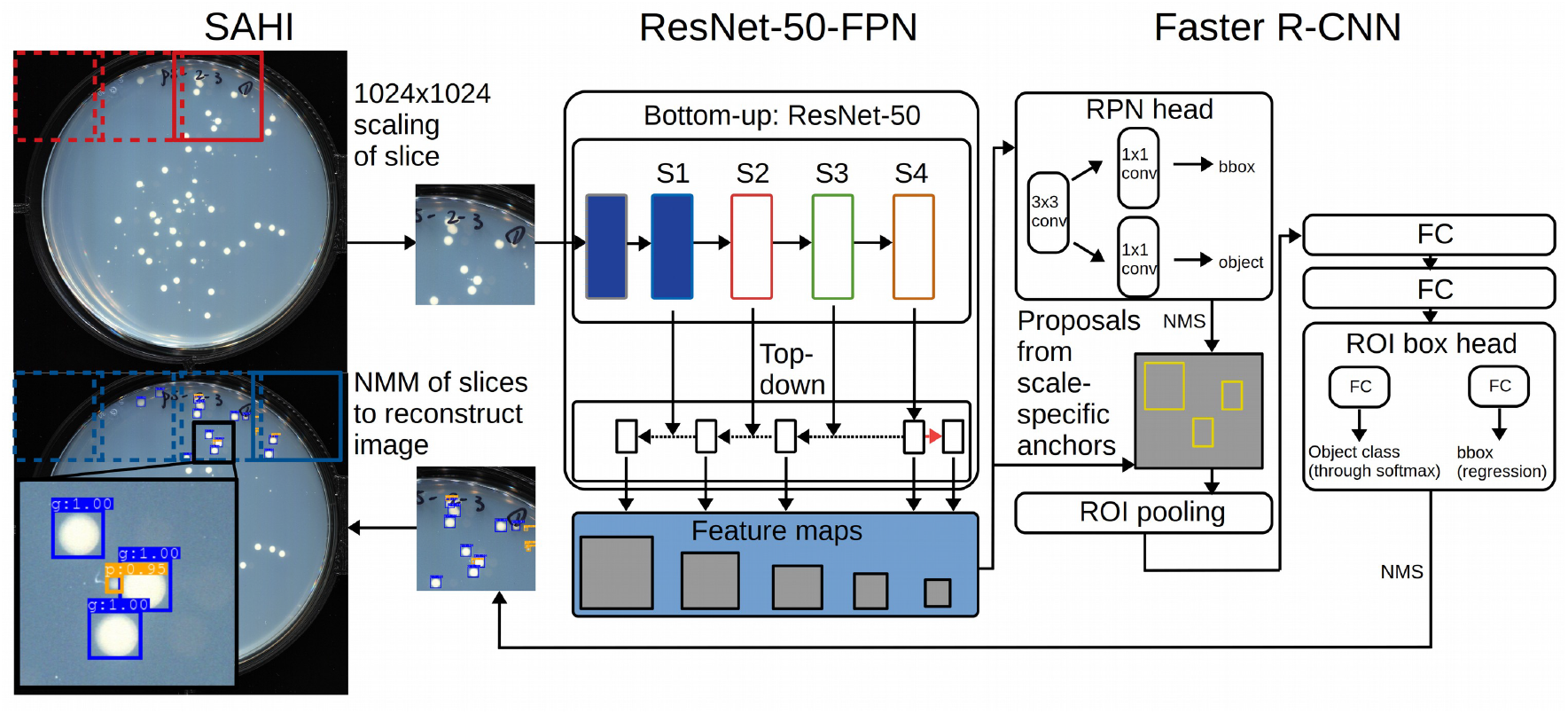
The architecture of the object detection pipeline. The entire plate image is first sliced into cropped images (red boxes) using a sliding window with the SAHI package. The image slice is scaled to 1024×1024 pixels regardless of its size and is passed into a ResNet-50-FPN convolutional neural network backbone. Here, Sx represents the convolutional stage of the ResNet-50 network. Solid blue indicates stages that were frozen during training. Convolution block outputs are rescaled (dotted arrow), undergo element-wise addition (intersection of solid and dotted arrows), and in one case, undergo max pooling (red arrow). This is the feature pyramid underlying the FPN (feature pyramid network) nomenclature. The output of this step is a collection of feature maps of various scales (resolutions) and semantic values. All feature maps are passed into the Faster R-CNN object detection architecture. In this architecture, the region proposal network (RPN), which is fully convolutional, generates region proposals that are likely to contain objects. These proposals are then passed to a final regressor and classifier which outputs a predicted bounding box and probability score for the class of the predicted object. Bounding boxes per image slice are filtered through non-maximal suppression (NMS). Finally, bounding box predictions on all image slices are merged into the final image using a non-maximum merging algorithm (NMM). The final output is a set of predicted bounding boxes on the entire image with associated Grande/Petite class probabilities (lower left image and zoomed portion).

### Model training

Network weights were initialized with a network pre-trained on the Microsoft Common Objects in Context (COCO) dataset (Lin et al., 2015) with 80 classification categories. Network weights were frozen from stage 1 to the beginning of the ResNet-50 backbone (indicated by solid blue blocks in Figure 2). The rest of the network was trained for 17 epochs with a batch size of 50 through stochastic gradient descent with a learning rate of 0.001, momentum 0.9, and weight decay of 0.0001. We used a learning rate scheduler with step-based decay, a linear warmup of 500 iterations, a warmup ratio of 0.001, and decay steps of factor 10 at 8 and 11 epochs. Images fed to the network during training were 512×512 px image crops of Petri dishes upsampled to 1024×1024 px. The crop size was selected so that objects in cropped images were of comparable size to objects from the COCO dataset. This step was important as frozen network weights were trained on objects of much larger relative size and correspond to these scale-specific features. Extensive data augmentation was also applied to training images, including random flips, 0.8 to 1.0 scale crops, and photometric distortions. These distortions included: random changes of brightness, contrast, saturation and hue, colour conversions between BGR and HSV, and randomly swapping colour channels (see details in MMDetection documentation; Wang, 2019).

The training parameters (including learning policies and optimizers) were selected following an extensive hyperparameter grid search on the validation image set. Once optimal hyperparameters had been selected for the network, we performed a hyperparameter grid search for SAHI. Bounding boxes were merged through NMM if they had a confidence score above 0.6 and had matching class predictions with an IOS of greater than 0.5.

### Model evaluation

During model finetuning, model performance was evaluated through the mean average precision metric with a bounding box intersection over union threshold (IOU) of 0.5 (mAP@0.5) on the validation set. Mean average precision is the area under the precision-recall curve across all bounding box confidence scores. Precision = TP/(TP +FP) and recall = TP/(TP+FN) with TP, FP, and FN, being true positives, false positives, and false negatives, respectively. For SAHI fine-tuning, we optimized parameters on the absolute error between predicted and ground truth Petite frequencies on images. However, this yielded similar parameters and performance to optimizing SAHI on <u>mAP@0.5.</u> For model performance on the test set, we report the mean average precision over IOUs ranging from 0.5 to 0.95 in 0.05 increments (mAP@0.5:0.95), alongside <u>mAP@0.5,</u> and <u>mAP@0.75,</u> which are commonly used in object detection competitions. We also show the Grande and Petite precision and recall. Finally, we compared predicted and ground truth Petite frequencies in the test set to evaluate real-world performance.

### Pipeline implementation

*petiteFinder* is implemented in Python 3.7.11 and uses the MMDetection package (Wang et al., 2019) to build, train, and test the model. It also uses SAHI (Akyon et al., 2022) which relies on MMDetection to perform sliced predictions. Before images are passed through the pipeline, slice heights and widths are determined by the image resolution equation, 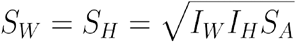, where *S*_*W*_, *S*_*H*_, *I*_*W*_, *I*_*H*_ are slice and image widths and heights, respectively. *S*_*A*_ is the relative area of the image slices to the area of the training images, which for our training set was 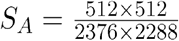. Assuming a user provides images of Petri dishes that fill the frame, this equation computes image slices with colony sizes relative to the background that are comparable to the training images. Training/testing was performed on an Nvidia GTX 1070 and GTX 1080 GPU with CUDA toolkit 11.3.1.

Users of *petiteFinder* have the option to output a CSV of Petite frequencies, visually annotated images with bounding boxes and scores, and a COCO formatted JSON file of annotations.

There is also an option to amend *petiteFinder* annotations with a GUI that enables users to explore predictions and add/remove bounding boxes around colonies (see Figure 3).

**Figure 3:**
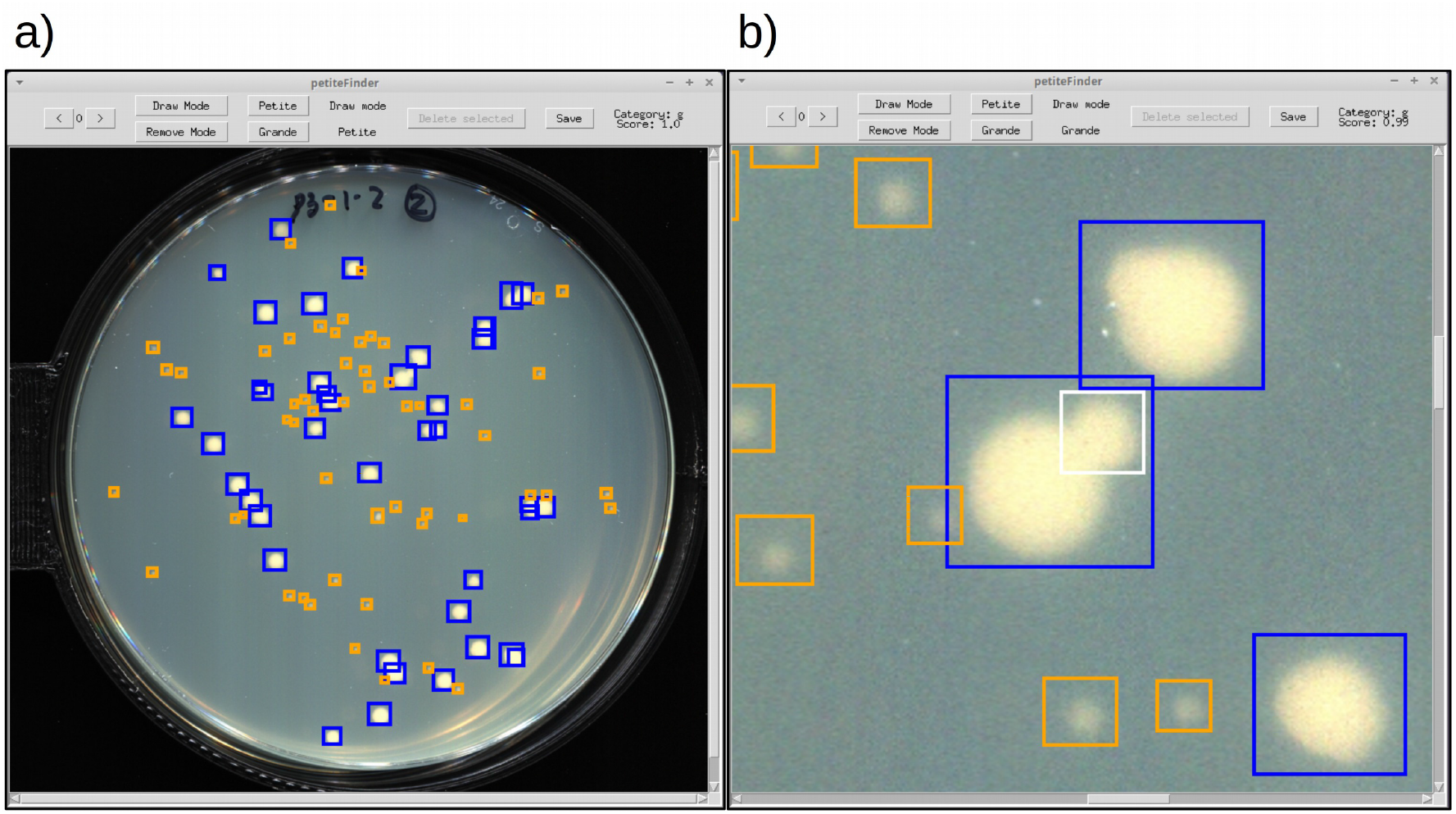
An example of the interface in the amend function within *petiteFinder*. a) The default view upon opening the GUI. Orange and blue boxes are Petites and Grandes, respectively. Arrow buttons in the top left can be used to move through images that have predictions. Hovering over colonies with a cursor reveals their class and probability score in the upper right. b) An example where a user is zooming into an image to add a new Grande bounding box that is being drawn in white in the center of the frame. Upon finishing the drawing movement with the mouse, the bounding box switches to the appropriate colour (blue) in this case. Functions also exist to remove bounding boxes after being selected with the cursor.

### Experimental details and best practices

#### Petri dish imaging

Six 100mm Petri dishes were scanned at a time, bottom-up, on an Epson V370 photo scanner at 600 DPI. The scanner was equipped with a custom 3D printed insert that dishes were placed into on the scanner surface and a black felt backing above the imaging surface. The 3D printed insert fixes Petri dish location for ease in cropping and reduces refraction from neighbouring Petri dishes which cause imaging artifacts. The black felt backing produced a dark background in scans which increased contrast between colonies and media in the images. Individual Petri dishes were cropped from the initial scan to fill the frame, resulting in 2376 × 2288 resolution images per petri dish. For best performance, images should be cropped so that Petri dishes fill the frame, as calculations of slice size are performed in the pipeline under this assumption. It is also recommended to have a resolution of at least 1000×1000 px per petri dish image to obtain comparable results to those reported in this study for similar sized colonies.

#### Media and growth conditions

Yeast colonies were grown on both complex and synthetic media to capture potential imaging variability in the most widely used growth media. The complex media was YPADG (1% yeast extract, 2% bacto-peptone, 0.1% glucose, 3% glycerol, 0.072% adenine hemisulfate). Synthetic media was SC-ura-trp (0.67% bacto yeast nitrogen base w/o amino acids, 0.1% glucose, 3% glycerol, 0.2% dropout powder lacking uracil and tryptophan). The shared characteristic across both media types that enables Petite identification is the carbon composition with reduced glucose which causes Petites to appear smaller and more translucent after 3-5 days of growth at 30°C (Dimitrov et al., 2009). Before plating, liquid cultures were grown at 30°C in a linear shaking water bath, while solid media growth took place in a forced air incubator at 30°C. Across the labeled dataset, images were taken in a range from 3-5 days after growth, resulting in varying relative sizes of Petite/Grande colonies. However, Petites were discernible by eye after this timeframe in all images.

#### Yeast strains

All strains used in Petite frequency experiments were in the W303 background. Yeast cultured on SC-ura-trp (0.1% glucose, 3% glycerol carbon source) were matings between W303 *MAT*a *leu*2-3,112 *can*1-100 *ura*3-1 *ade*2-1 *his*3-11,15 and W303 *MAT*α *leu*2-3,112 *can*1-100 *ade*2-1 *his*3-11,15 *trp*1-1. Yeast cultured on YPADG was a haploid strain W303 *MAT*a *leu*2-3,112 *can*1-100 *ura*3-1 *ade*2-1 *his*3-11,15.

## Results

### Pipeline performance

To evaluate the performance of the pipeline, *petiteFinder* was applied to the images in the test set and compared against ground truth annotations. The test set consists of 17 petri dish images (4 YPADG, 4 SC-ura-trp nonideal, 9 SC-ura-trp ideal) with 1327 ground truth colonies. Of these 1327 colonies, 602 are Grande and 725 are Petite. On the test set, *petiteFinder* achieved a mean average precision at IOU=0.5 of 0.96 with precision and recall of 0.96 and 0.99, respectively, for both Grandes and Petites (Table 1). These results are without any post-processing or user input. To improve accuracy even further, users can cull model predictions below a particular quality score.

**Table 1:** Ground truth annotations of 1327 colonies were compared to predicted annotations from *petiteFinder* across 17 images in the test set. The top row includes mean average precision computations (mAP) across IOU thresholds from 0.5 to 0.95 in 0.05 increments, at 0.5 IOU, and 0.75 IOU. Below this, precision TP/(TP + FP) and recall TP/(TP+FN) at 0.5 IOU have been computed for each colony class.

| mAP(0.5:0.95): 0.64 |  |  | mAP@0.5: 0.96 |  |  | mAP@0.75: 0.62 |
| --- | --- | --- | --- | --- | --- | --- |
| Category |  |  | Precision |  |  | Recall |
| Grande |  |  | 0.96 |  |  | 0.99 |
| Petite |  |  | 0.96 |  |  | 0.99 |

To test what these accuracy metrics mean for computations based on colony detections, we compared Petite frequencies determined by *petiteFinder* and manual counting for each petri dish (Figure 4a). The red points are predictions, and the black points are manual counting (ground truth) Petite frequencies. Across all test images, the average absolute difference in predicted vs. ground truth Petite percentages (prediction error) is 1.7%, with a standard deviation of 1.2%. To understand the potential impact of this error on biological insights gained from *petiteFinder* predictions, we compare the prediction error to expected variation in Petite frequencies from experimental sampling. Assuming that the generation of Petite colonies is a Bernoulli process, the standard deviation in Petite frequency expected from sampling numerous Petri dishes with the same underlying Petite probability is 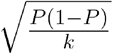. Here *P* represents the probability of Petite colony production and *k* is the number of colonies per Petri dish. Taking *p* to be the ground truth Petite frequency, and *k* = 78, which is the average number of colonies per plate in the test set, we plot this binomial sampling error as a gray envelope around the ground truth measurements with +/- the binomial standard deviation (gray envelope in Figure 4a). Above Petite frequencies of 0.1, Figure 4a demonstrates that *petiteFinder* predictions are within this sampling error envelope, suggesting that prediction errors are less impactful than sampling error itself. Below 0.1 petite frequency, where sampling error is negligible, predictions exist outside this envelope but still yield small absolute error compared to ground truth measurements (<5% error in Petite percentage).

**Figure 4:**
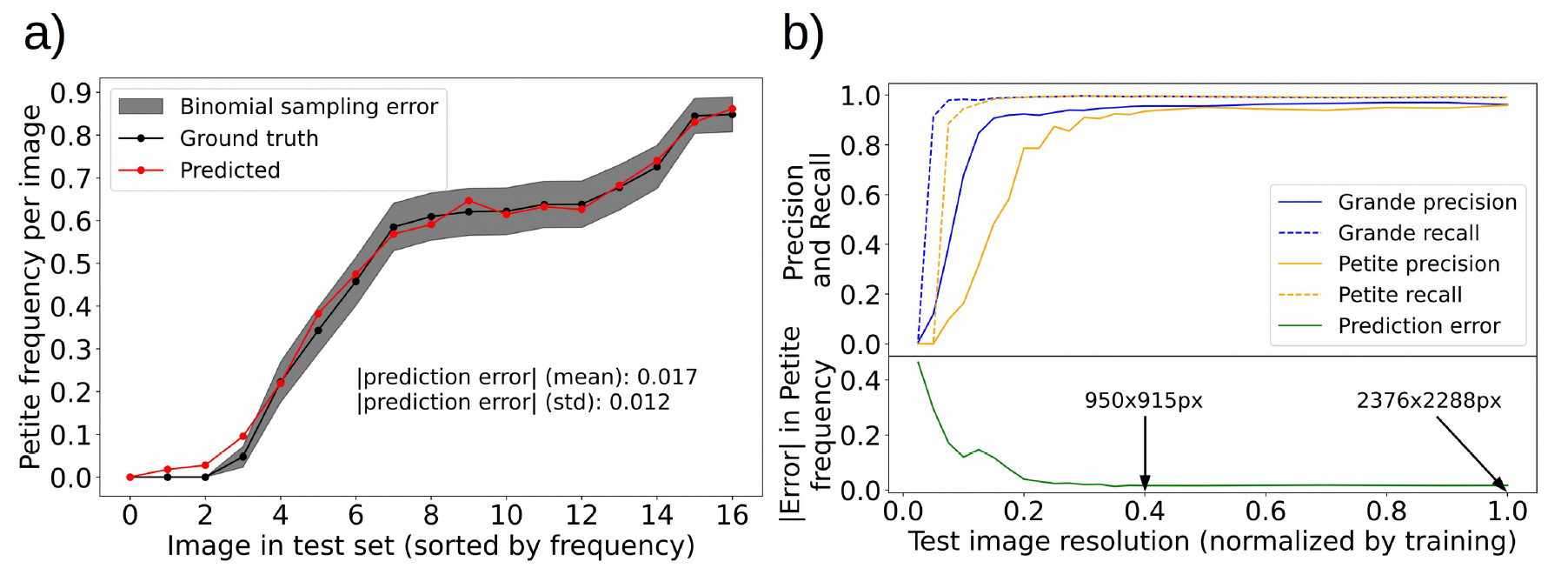
*petiteFinder* performance compared to manual counting alongside a test of model robustness. A plate-wise comparison of predicted and manually counted Petite colony frequencies. The red curves are the predictions and the black curves are manual counting. The average absolute deviation between predicted and ground truth Petite percentage (prediction error) is 1.7%. The gray envelope is the binomial sampling error (ground truth +/- standard deviation), assuming that Petite production is a Bernoulli process with a probability equal to the ground truth frequency when sampling 78 colonies per plate image. Top panel: Precision and recall of each colony category as a function of test image resolution normalized by the image resolution of the training data. Bottom panel: Absolute error in petite frequency as a function of test image resolution. The error is defined as the deviation between predicted and ground truth Petite frequency. Labels are also included to denote absolute image resolutions.

While we applied extensive data augmentation during training that would encompass a variety of imaging conditions, one parameter that remained fixed in both training and testing was the image resolution. Therefore, to understand how robust *petiteFinder* is to image resolution, we plotted model accuracy metrics while varying the image resolution of the test set in Figure 4b. The x-axis in this plot should be interpreted as the fractional size in x and y dimensions of the test image relative to the training image resolution (2376 × 2288 px). Recall of the model in the top panel of Figure 4b reveals that below 0.1 test image resolution (238 × 229 px) all colonies disappear according to the model due to their small size. As expected, this occurs first for Petites which are composed of fewer pixels. As we move towards lower resolutions starting at ∼0.4 of test image resolution (950 × 915 px), we see that Petite colony precision drastically decreases due to misclassifications of small imaging artifacts as Petites. The same decrease in precision occurs for Grandes but at even lower test image resolutions. Petite frequency prediction error is also shown in the bottom panel as a function of test image resolution. Remarkably, at 0.4 test image resolution and above (950 × 915 px or ∼1M px), prediction accuracy is comparable to the results on our 2376 × 2288 px resolution plate images. In general, the resolution threshold where Petite detection degrades most rapidly is dependent on experimental conditions that dictate Petite colony size. For our particular experimental conditions, this threshold indicates that a rapid degradation in Petite detection performance occurs below 4x the resolution where Petite colonies vanish.

## Discussion

In this study we introduced *petiteFinder*, a computer vision tool to detect Grande and Petite colonies and determine Petite colony frequencies from images of Petri dishes. *petiteFinder* is the first computer vision pipeline tailored for this application to the best of our knowledge. We showed that it performs comparably to human annotation, with errors that are smaller on average than the theoretical sampling error. Overall, the average difference in computed Petite percentages between *petiteFinder* predictions and manually annotated colonies was 1.7%. We also detailed best practices for using the tool, including minimum image resolutions and protocols to recreate imaging conditions. When *petiteFinder* is run on a GPU such as an Nvidia GTX 1070, each Petri dish image is processed in ∼2 seconds. Depending on colony density, this is a 10-100 fold improvement in throughput relative to manual counting without any user input. *petiteFinder* can also be run on a CPU, and, although the inference will be significantly slower (∼2 minutes per image), it still comes with the benefit of being completely automated. We believe that this advancement in throughput and detailed protocols for imaging and experimental conditions will drastically increase this assay’s statistical power and repeatability.

While on average *petiteFinder* performed admirably, under 0.1 Petite frequency we see a larger prediction error than binomial sampling error from a distribution of cells with a fixed probability of becoming Petite. This is due to the sensitivity to false positives at low Petite frequencies. To address this issue and enable modifications to model predictions, we designed a GUI that allows experimentalists to visually explore and amend prediction outputs from *petiteFinder*. This ‘amend’ function lets users draw and remove bounding boxes around colonies following the initial prediction. Beyond performance and usability, we also emphasized how this computer vision problem of identifying Petite colonies highlighted ongoing difficulties in small object detection. We motivated changes to existing object detection architectures that were necessary to detect Petite colonies in this study, even with state-of-the-art architectures.

Regarding the development of this pipeline, a natural question is whether or not a supervised deep learning architecture is necessary to solve this problem with sufficient accuracy. Numerous popular semi-supervised approaches exist for colony segmentation, such as CellProfiler (Bray et al., 2015) and AutoCellSeg (Khan et al., 2018). These pipelines use a variety of thresholding and edge detection algorithms to segment colonies from the background of plate images. However, they require user input on plate edge detection in the case of CellProfiler, and user input on example colony locations in AutoCellSeg. Nevertheless, at the segmentation task, these approaches perform admirably. For example, AutoCellSeg achieves a segmentation accuracy of 97% (Khan et al., 2018). Given segmentations from these types of approaches, users can apply a variety of classification methods to categorize colonies based on their features.

We initially took this same type of semi-supervised approach when exploring the Petite colony detection problem, as detailed in the supplementary information of this study (see an overview of data and approach in Figure S1 and S3). Instead of modifying existing approaches for colony segmentation, we created our semi-supervised pipeline which incorporates Otsu’s thresholding (Otsu, 1979), watershed segmentation (Digabel & Lantuejoul, 1978), and plate detection with a Hough transform (Hough, 1962). We then relied on two features of Petite/Grande colonies that were biologically relevant to the classification problem, the size and intensity of colonies, and clustered colonies based on these features with average-linkage agglomerative clustering. Our unsupervised clustering approach on human-annotated bounding boxes yielded a Petite precision and recall of 0.93 and 1.00, respectively, and Grande precision and recall of 1.00 and 0.91, respectively. However, the difficulty was that even with perfect segmentation, the clustering resulted in exceptionally poor Petite frequency predictions in some images (Figure S2). Beyond reduced classification accuracy, we also required numerous user inputs in the full semi-supervised pipeline, as shown in Figure S4. At its best, the semi-supervised pipeline including segmentation and classification yielded a Petite precision of 0.75 and recall of 0.87. It also achieved Grande precision and recall of 0.98 and 0.68, respectively. These results were obtained for one media type under ideal imaging conditions (Figure S5 and supplementary Table S1). This relatively poor predictive capacity strongly suggested that with more media types and variations in imaging conditions, this would not be a robust model useful to researchers. This negative result spurred the development of the supervised pipeline in this study.

Finally, it is important to comment on comparisons to existing colony detection methods. Even with perfect colony segmentation, we show in Figure S2c and S2d that the unsupervised classification we explored (clustering colony size and intensity) will perform more poorly than *petiteFinder*. As it stands, robust colony segmentation also requires user input in existing semi-supervised pipelines. For this reason we believe an exhaustive search of modifications to existing semi-supervised pipelines is unwarranted. There is, however, a supervised red/white colony detection pipeline described in (Carl et al., 2020) that can be used for Petite frequency calculations in strains with an appropriate genetic background. In yeast strains with specific mutations in the adenine synthesis pathway, an intermediate compound (initially white) accumulates and turns red when oxidized in respiring cells. It is also possible to treat cells directly with tetrazolium to induce a rapid transition from white to red, specifically in Grande cells (Hess et al., 2009). While the pipeline in (Carl et al., 2020) performs admirably, and could be modified for Petite detection with the red/white colony assay, there are two disadvantages to using the red/white assay over our experimental approach. First, the red/white assay requires experimental constraints on media (adenine limited media) and the genetic background of the strains, whereas large/small colony detection only requires constraints on the media. Second, for red pigments to form, experimentalists often need to wait further after colony growth or apply chemical treatments which decrease experimental efficiency compared to our approach. With respect to the object detection pipeline in (Carl et al., 2020), one downside is that it requires post-processing of segmentation results, including heuristics on eccentricity and absolute size of colonies in pixels. Furthermore, assuming perfect segmentation in the study by Carl et al. (where segmentation accuracy is not reported), *petiteFinder* still achieves higher accuracy in Grande/Petite classification compared to white/red classification in experiments.

## Conclusion

In this study we developed *petiteFinder*, an automated computer vision tool to detect Grande and Petite yeast colonies and determine Petite frequencies from images of Petri dishes. Colony detection with *petiteFinder* results in high accuracy Petite and Grande localization in images in a completely automated fashion. It achieves accuracy comparable to human annotation but at up to 100 times the speed and outperforms semi-supervised Grande/Petite colony classification approaches. By constructing this tool and providing details of experimental conditions, we hope this study will enable larger-scale experiments that rely on Petite colony frequencies to infer mitochondrial function in yeast.

## Data availability

Source code, detailed documentation on installation and use, modifiable 3D print files, and all labeled data in this study are available at www.github.com/javathejhut/petiteFinder.

## Author contributions

C.J.N conceptualized the project. Both C.J.N. and E.K. implemented, modified, and trained computer vision models in hyperparameter grid searches. C.J.N. and E.K. also wrote the software for both the GUI and CLI. E.K. wrote tool installation and usage documentation. C.J.N. performed the experiments and labeled data, evaluated model performance, created manuscript figures, and wrote the initial draft. E.K. edited the manuscript. C.J.N constructed and evaluated the semi-supervised approach in the SI.

