## Supplementary information for "petiteFinder: An automated computer vision tool to compute Petite colony frequencies in baker’s yeast"

Christopher J. Nunn[1] and Eugene Klyshko[1,2]
[1]Department of Physics, University of Toronto, Ontario, Canada, M5S 1A7
[2]Department of Chemical and Physical Sciences, University of Toronto Mississauga, Ontario, Canada, L5L 1C6

**Supplementary Information:** A semi-supervised Grande/Petite colony detection approach as a stepping stone towards *petiteFinder.*

### Description of the labeled dataset (smaller and fewer media types than *petiteFinder)*

The dataset used in this project is a collection of 59 petri dish images of yeast colonies on synthetic dropout media with glucose (0.1%) and glycerol (3%) as carbon sources. Mated Grande and Petite colonies were plated onto this media, with Petite progeny showing up as smaller less opaque colonies compared to their Grande counterparts after 3-5 days of incubation at 30$^{o}$C. Six plates at a time were scanned after the same period of growth on a computer scanner. They were then cropped to images containing an individual plate with the aid of a 3D printed scanner insert that fixed their positions and also reduced refraction due to adjacent plates. These images were annotated using the LabelMe package (Russell et al., 2008), where bounding boxes were drawn around Grande and Petite colonies and assigned to their corresponding labels. In total 4868 bounding boxes are in this dataset, with 2684 Petite and 2184 Grande labels. Variation in agar concentrations within the media poured into plates in this dataset produced 20 images we consider "non-ideal" due to the diffuseness of colonies, and 39 images we consider "ideal". For the purposes of testing the predictive capacity of an unsupervised object classification pipeline we separated "ideal" and "non-ideal" images into 80% validation sets and 20% test sets. See Figure S1 for a summary of this dataset.

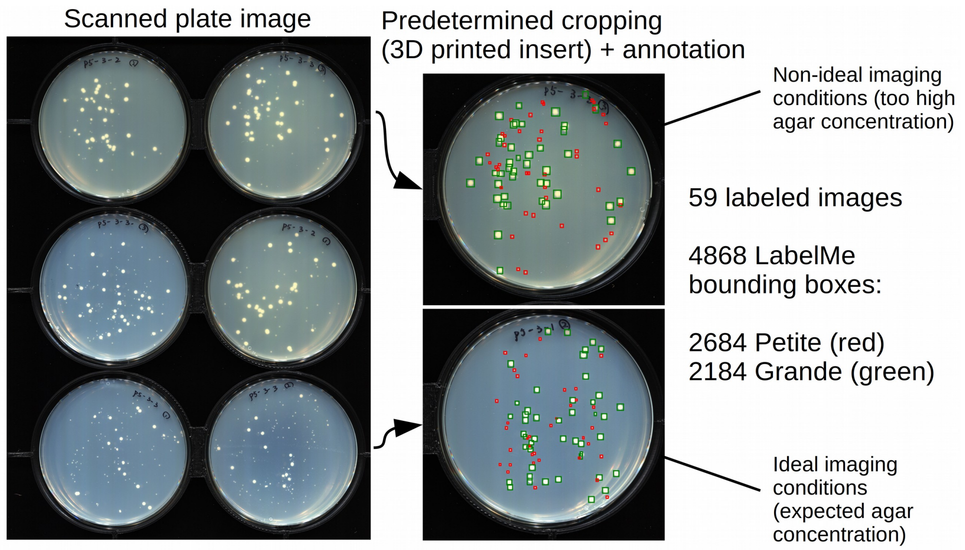


**Figure S1:** An overview of the labeled dataset for the semi-supervised approach. Petri dishes of synthetic complete media and Grande/Petite yeast colonies were placed into a 3D printed insert and scanned on a computer scanner bottom-up (leftmost image). Individual plate images were cropped out of the large image, and 59 of these images were annotated using the LabelMe annotation tool (Russell et al., 2008). Grande and Petite colonies are indicated by green and red bounding boxes, respectively. Experimental variation in media preparation/pouring of plates produces diffuse low quality scans and sharper plate images as shown by the upper and lower plate in the middle panel.

### An unsupervised Petite colony detection approach

#### *Feature selection and clustering of labeled data*

We built the core of the unsupervised classification algorithm by selecting distinguishing features of Grande/Petite colonies expected from biological understanding. Petite colonies are smaller than Grande colonies due to the inability of Petite cells to switch to respiration once carbon sources are consumed. This also means that the density of cells in Petite colonies should be lower than Grandes because respiring cells are unable to grow on top of the region of consumed glucose. This suggests that differences in both the size of Grande and Petite colonies and their pixel intensities might enable their classification. To test this idea, we took all 59 annotated images in the labeled dataset and clustered the annotated bounding boxes surrounding Grandes and Petites in size and intensity (Figure S2). Figure S2a shows an example plate image with bounding boxes around colonies. Figure S2b shows the average-linkage hierarchical clustering with Euclidean distance of these bounding boxes in size and intensity, where clusters with a larger average size are considered Grandes. This clustering method was selected because there was no implicit assumption of equal cluster sizes, and it was sufficiently sensitive to outliers which is important for plate images with low/high Petite frequencies.

We evaluate the effectiveness of this classification scheme on all 59 plate images by considering the precision and recall of Petite classification (Figure S2c). The precision is $\frac{TP}{TP+FP}$, where $TP$ are the true positive classification counts and $FP$ are the false positive counts. The recall is $\frac{TP}{TP+FN}$, where FN stands for the false negative counts. The precision and recall for Petites across all 59 images is 0.933 and 0.999, respectively. The Grande precision and recall is 0.999 and 0.912, respectively. These metrics suggest that the classification scheme struggles with false positives Petite classifications, which are likely due to imaging artifacts. It also struggles to identify all Grande colonies in images, which likely occurs when Grande colonies are merged together when plated at high density. In Figure S2d we show the Petite frequencies predicted per plate according to this scheme compared to the ground truth from annotated data. It highlights that while we have good average Petite classification accuracy (average deviation of 0.038 in Petite frequency, standard deviation of 0.058), plates where there are a small number of colonies or low Petite frequencies can produce significant deviations from ground truth measurements. Overall, however, these results were promising and suggested that this type of classification scheme combined with a colony detection pipeline may perform reasonably well. We discuss the construction of this joint colony detection and classification pipeline in the next section.

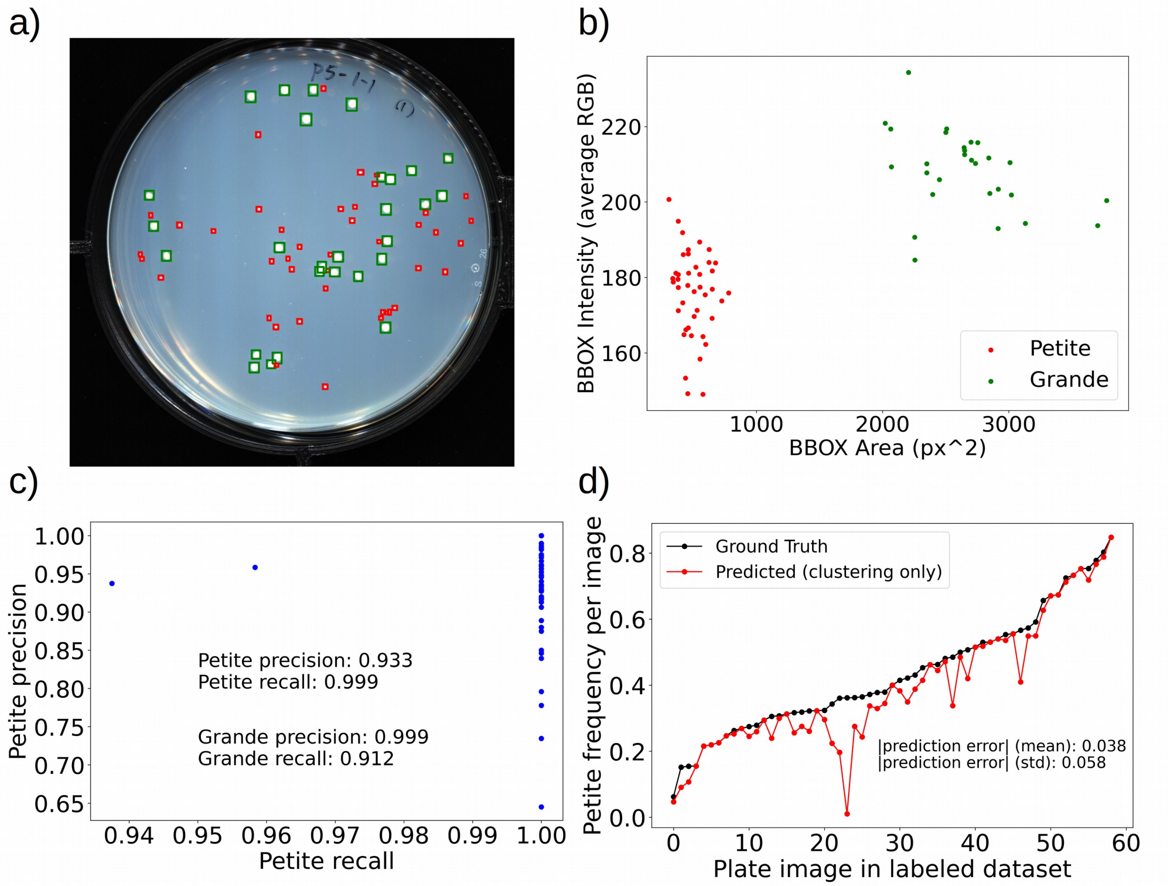


**Figure S2:** Performance of unsupervised average-linkage hierarchical clustering on the bounding boxes of labeled data. a) An example plate image with annotated bounding boxes around colonies generated through the LabelMe annotation tool. b) Average-linkage clustering result of bounding boxes (BBOX) in area and average intensity of the contained pixels from the image in (a). c) Precision and recall of this unsupervised classification of Petite bounding boxes across all 59 labeled images. Each dot represents the precision and recall of each plate image for Petite classification. Also noted are the precision and recall for Grandes which are not shown. d) The Petite frequencies computed per plate in the labeled data using this clustering approach as well as the ground truth petite frequency determined through human annotation. Average and standard deviation in prediction error is also denoted, where prediction error is the difference between ground truth and predicted petite frequencies per image.

*Combining colony detection and classification*

In order to create an automated computer vision pipeline that accepts plate images and computes Petite frequencies we needed to combine methods to detect colonies and the previously explored classification scheme. To detect colonies, we opted to use a thresholding approach to separate colonies which are brighter in images from their darker uniform background on the agar surface (Figure S3).

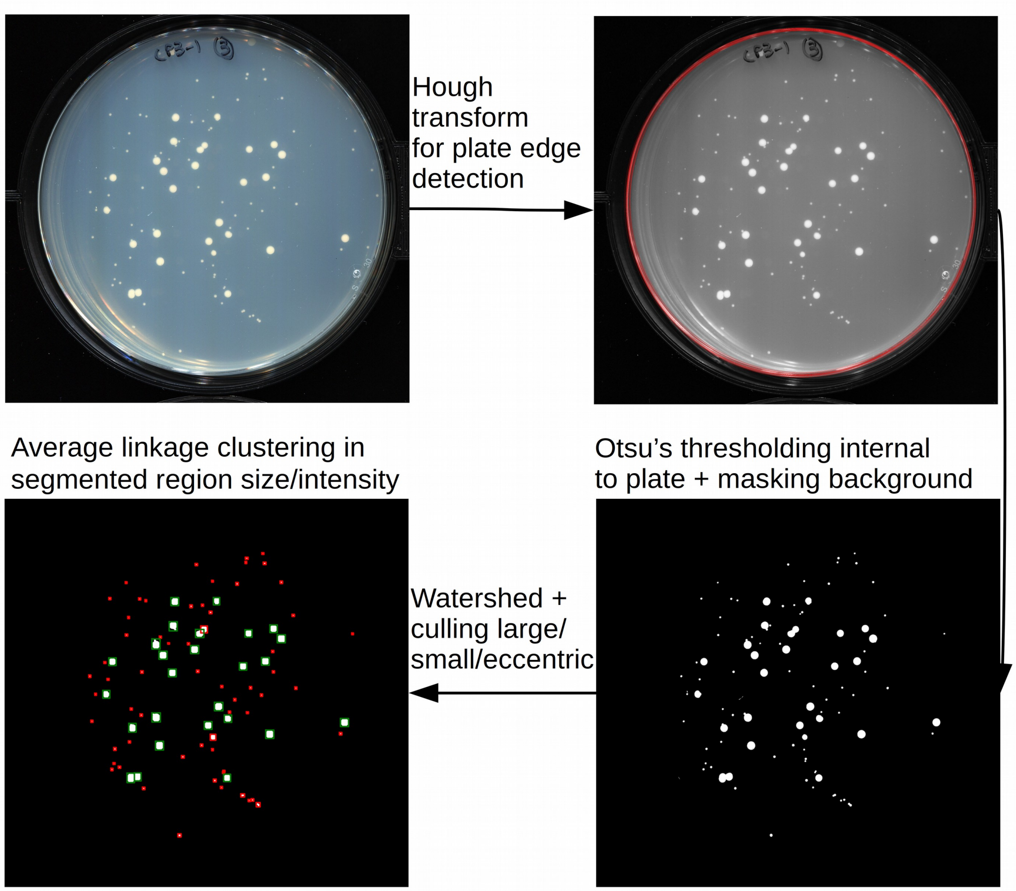


**Figure S3:** An overview of an unsupervised automated colony classification pipeline that identifies colonies and classifies them as Petite or Grande based on the clustering approach detailed in Figure S2. A Hough transform accepts a user suggested radius parameter (to account for various plate and image sizes) and outputs the predicted plate boundary in red. A mask is applied to the exterior of the plate, followed by Otsu's thresholding to segment the image into foreground (white), and background (black). Given a maximum ($args.max$) and minimum ($args.min$) expected colony size provided by the user, a watershed transform is applied to the foreground with a minimum displacement between basins of $\sqrt{\frac{args.max}{4\pi}}$ to separate only the merging of large colonies which is most common. Colonies with size above $args.max$, below $args.min$, and with eccentricity >0.9 are removed. Finally, the labeled foreground objects are clustered using average-linkage clustering in size and intensity features of the labeled region.

To start, we apply a Hough transform (Hough, 1962) to identify the plate edge with an estimated plate radii range provided by the user. An example plate edge detection is shown in red in the upper right panel of Figure S3. This allows us to mask out the exterior of the plate and restrict the colony detection to real ones on the agar surface. This is followed by Otsu's thresholding (Otsu, 1979), which is an algorithm that selects a foreground/background intensity threshold by finding the value that minimizes the intensity variation within each class. Given maximum ($args.max$) and minimum ($args.min$) expected colony sizes provided by the user, we then apply a watershed transform (Digabel & Lantuejoul, 1978) with a minimum separation between minima of watershed basins of $\sqrt{\frac{args.max}{4\pi}}$ to focus on separating merged Grande colonies. This is followed by the removal of colonies with sizes above $args.max$, below $args.min$, and with eccentricity >0.9. Filtering out only high eccentricity objects we found removes fabric fibres from images, but retains segmented merged colonies that don't necessarily have an eccentricity near zero. Finally, segmented regions are clustered according to the previous section assuming that they are colonies.

*Evaluation of the combined colony detection and classification pipeline*

To evaluate how effective this complete Petite classification pipeline is we must first address three important points:

The first is that $args.min$ and $args.max$, the expected minimum and maximum colony sizes, must be selected for the set of data we are testing this pipeline on. We do this by testing numerous ranges of these parameters for the "non-ideal" and "ideal" validation image sets and picking the parameter values that maximize performance by computing the $F_1$ score $(\frac{2*precision*recall}{precision+recall})$. The maximum value of this score indicates the best result across all parameter regimes if precision and recall are equally weighted. With this approach the pipeline becomes semi-supervised, which is similar to users inputting these maximum and minimum colony values per image or batch of images and is common in modern colony detection approaches (Khan et al., 2018; Carl et al., 2020).

The second point is that to compare predicted bounding boxes (the rectangles that bound unique segmented regions from the pipeline), and ground truth bounding boxes from human labeling, we have to consider the overlap of these rectangles. The most common approach is to compute the intersection area over union (IOU) of the predicted and ground truth bounding boxes and provide an IOU threshold above which we consider a good prediction. As will be discussed later, the thresholding approach we took in the pipeline often reduces the size of Petite colonies due to their transparency and therefore similarity to the background. Thus, we choose an IOU threshold of 0.1 above which we consider bounding boxes to overlap to account for the shrinking of Petite colonies after thresholding.

The third point is that to really evaluate the accuracy of this now semi-supervised approach we individually tune colony size parameters in the validation image sets, which is similar to users selecting maximum/minimum sizes per image, but test the predictive value of the pipeline on separate test images.

As a demonstration of the colony size parameter tuning, we show the precision and recall for both Grande and Petite colonies in the "ideal" validation image set (31 images) in Figure S4 across a variety of minimum/maximum colony sizes. In the computation of recall and precision in this figure we use an IOU threshold of 0.1. In Figure S4a and S4b we can see that the optimal maximum and minimum colony area values of 4000px and 20px provide an average precision and recall for the detection of Grandes of 0.98 and 0.73 respectively.  In Figure S4c and S4d we see that the best average precision and recall for Petites are 0.73 and 0.8, respectively.

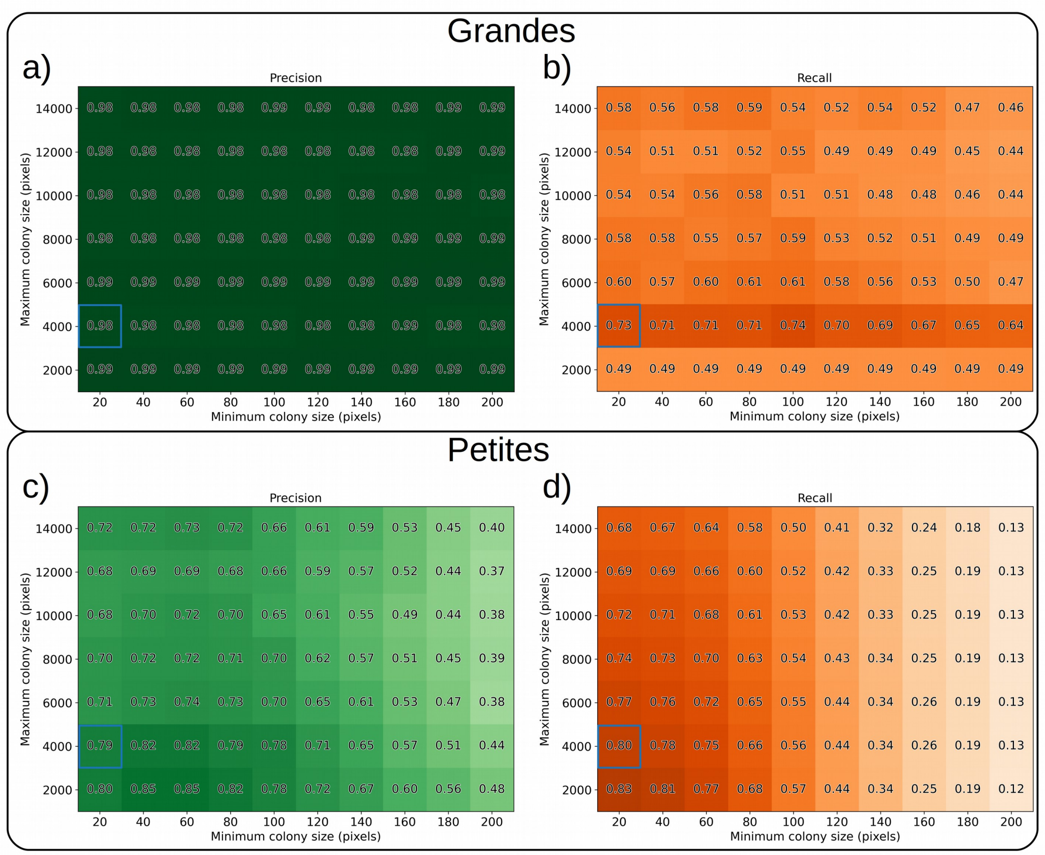


**Figure S4:** Precision and recall of the unsupervised pipeline for both categories across numerous $args.min$ and $args.max$ parameters for the “ideal” images in the labeled validation set. The parameter regime that maximizes the $F_1$ score $\left(\frac{2*precision*recall}{precision+recall}\right)$ averaged across each category is highlighted in blue.

Given a prescription for how to choose minimum/maximum colony sizes to optimize prediction we now test how predictive the semi-supervised pipeline is for the “ideal” test image dataset (8 images) in Figure S5. In Figure S5a we show the precision and recall for Petite and Grande classification as a function of IOU. We can see clearly that Petite colonie classifications are more sensitive to IOU thresholds because of the similarity of their intensities to the background intensity. We see reasonably high precision and recall for Petites (~0.8) at an IOU of 0.1, but a Grande recall near 0.6. This is due to either the elimination of Grandes according to the maximum colony size selected or Grandes being classified as Petites. In Figure S5b we also [illegible]

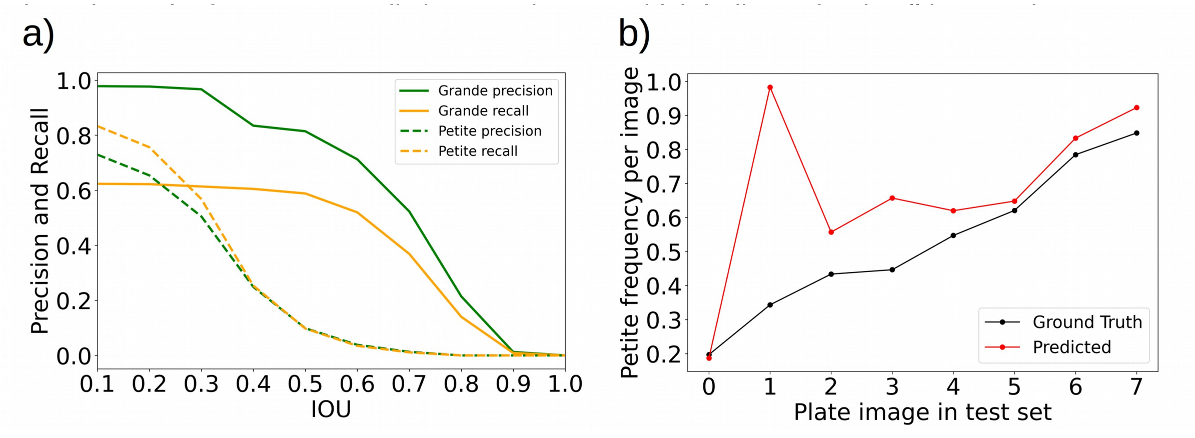

over union (IOU) of the predicted and ground truth bounding boxes in the “ideal” test dataset. Here the min/max colony size parameters are those highlighted by the blue boxes in Figure S4. b) Petite frequency predicted by the automated unsupervised pipeline compared to the ground truth human annotations (test set).

We summarize the effectiveness of this colony detection pipeline in Table 1, where we include optimal maximum/minimum colony sizes inferred from the “ideal” and “non-ideal” image validation sets and precision and recall in the corresponding test sets. As can be seen from this table, the optimal colony size thresholds vary significantly due to experimental variation and so does the classification accuracy of this approach. This suggests that while we see reasonable Petite/Grande classification especially in the “ideal” test dataset using our approach, parameters like minimum and maximum sizes really do need to be tuned per image or batch of images to obtain the best performance if we are relying heavily on size as a feature. This can be remedied by an automated deep learning object detection approach, which for example can easily learn other defining features of colonies vs. noise or dust such as nonlinear radial intensity profiles and irregular colony boundaries.

| **Image test set (IOU=0.1)** | **Grande precision** | **Grande recall** | **Petite precision** | **Petite recall** | **optimal minimum colony size from validation set (pixels)** | **optimal maximum colony size from validation set (pixels)** |
|---|---|---|---|---|---|---|
| “Ideal” (8 plates) | 0.98 | 0.68 | 0.75 | 0.87 | 20 | 4000 |
| “Non-ideal” (4 plates) | 1.0 | 0.69 | 0.66 | 0.21 | 60 | 8000 |

**Table S1:** Precision/recall for both Grandes and Petites in the “ideal” and “non-ideal” test datasets.

**References:** Shared with the list found in *petiteFinder* maintext.
